# MIEN: a deep-learning model for pocket-conditioned glycan pose and conformation discrimination

**DOI:** 10.64898/2026.10.13.758804

**Authors:** Xirui Huang, Zhixin Tian

**Affiliations:** School of Chemical Science & Engineering, Shanghai Key Laboratory of Chemical Assessment and Sustainability, Tongji University, Shanghai 200092, China

**Keywords:** protein-carbohydrate interaction, glycan conformation, post-docking reranking, equivariant graph neural network

## Abstract

`In` protein-carbohydrate interaction, Molecular docking can generate many glycan poses that appear plausible, but the top-ranked pose may differ from the experimentally observed conformation. We introduce Minimal Interface Equivariant Network (MIEN), a structure-based equivariant graph network that scores the compatibility of a glycan pose with a fixed protein pocket. The model operates on a minimal-interface graph that contains only protein atoms forming polar or CH-π contacts with the glycan; protein and sugar residue names are excluded. MIEN was trained on 5,241 experimental complexes and one unified set of contact-preserving rigid-body negatives grouped into three semantic-contact tiers. On a protein-disjoint benchmark of 260 held-out complexes, MIEN separated experimental poses from non-native poses and ranked experimental poses more highly within matched pockets. We then evaluated MIEN on AutoDock-GPU and GlycanDock candidate pools generated independently for the same complexes. Across three training seeds, MIEN selected lower-RMSD top-1 poses more often than either docking engine and increased the number of best poses below 1, 2 and 4 Å. These results provide a practical route to more realistic protein-carbohydrate complex poses and a stronger basis for structural and mechanistic studies of protein-carbohydrate interactions.

## 1. Introduction

Protein-carbohydrate recognition takes part in cell-cell recognition, immune surveillance, host-pathogen attachment and metabolic regulation ^1,2^. Its physical chemistry is different from protein-small-molecule recognition, and this difference affects prediction. Protein-carbohydrate interfaces are often shallow and polar; hydrogen bonding and CH-π stacking are major interactions ^3,4^, and affinities are often in the high-micromolar to millimolar range ^5,6^. For this reason, many glycan placements can look stereochemically reasonable. The important biological question is whether a proposed pose matches the conformation and the local interaction geometry that are seen in an experimental complex.

Docking, structure prediction and protein design answer this question in an indirect way, because they generate candidate poses. One run can return many poses with similar docking scores, but the glycan placements can be very different. The scoring function does not always point to the pose that is closest to the experimental state ^5,7,8^. Audits of physical validity also show that a pose can look reasonable and still fail basic geometric or chemical checks ^9^. The selection problem is therefore different from binding-site detection and from cross-sugar classification; in this problem, the glycan and pocket are fixed, and the model has to decide which conformational state fits the local protein environment. A useful signal should therefore come from pocket-conditioned interaction geometry, not from the protein family or the glycan name.

Computational studies have not fully separated this decision from nearby tasks. Binding-site and pocket predictors show where a carbohydrate may bind ^10,11,12,13,14^. A recent deep-learning model in this group combines protein-language-model embeddings with structural features to locate carbohydrate-binding sites ^13^. Protein-glycan affinity models provide related information; CSM-carbohydrate predicts binding affinity and also gives a docking score function ^15^, and a structural-feature regression study linked contact potentials, interaction energies, binding-site composition and atom-level contacts to protein-carbohydrate affinity in several complex categories ^16^. DIONYSUS collects experimental protein-carbohydrate interfaces for large-scale analysis ^17^. Glycan-aware generative models have started to predict carbohydrate and glycoprotein structures directly ^18,19^. Complex-quality models test whether a proposed complex is native-like, but their benchmarks often mix placement errors, internal glycan conformation and ligand chemistry ^5^. Protein-level classifiers separate binders from curated non-binders, but they do not resolve the conformation inside a pocket ^20^. CLIMBS is an interaction classifier that distinguishes carbohydrate-protein interactions from synthetic negatives, whereas MIEN is a conformation reranker for candidate glycan poses inside a fixed pocket ^21^. Quantitative glycan-array models usually treat the glycan as a global ligand descriptor ^6,22,23^; these tasks are related, but none of them directly reports whether a given glycan geometry is compatible with an experimental complex.

Several docking and structure-modeling methods have been developed for glycan systems. Vina-Carb adjusts glycosidic angles during docking ^24^, GlycanDock refines protein-glycoligand poses in Rosetta ^25^, GlycoTorch Vina targets sulfated glycans ^26^, HADDOCK supports integrative protein-glycan modeling ^27^, AlphaFold 3 can model glycan-containing complexes but has limits in glycan stereochemistry and conformational sampling ^28^, and GlycoShape provides conformational ensembles for glycoprotein modeling ^29^. Related studies have examined atom-level protein-glycan interactions ^30^. Keys, Kiessling and coworkers analyzed the energetic landscape of CH-π interactions in protein–carbohydrate binding and showed that these contacts can make important contributions to recognition, with their strength shaped by local ring geometry and chemical environment ^31^. These studies also include carbohydrate simulations ^32^ and Rosetta glycan modeling ^33^. Together, these studies provide complementary structural and conformational context and motivate MIEN to treat CH-π and polar contacts as explicit semantic edges rather than generic distance-based contact pairs. Water-guided docking uses crystallographic water sites to improve glycan pose generation ^34^, machine learning has been used to select native-like protein-GAG poses ^35^, and pose-reranking models have been developed for small-molecule ensembles ^36^ and RNA-ligand docking ^37^. Structure-generation and validation studies also show that carbohydrate models can retain stereochemical or geometric errors ^38,39^. These studies address candidate generation, energy scoring or native-like complex assessment, but the problem of isolation of the fixed-pocket, fixed-glycan pose-ranking needs further investigation.

To learn this decision, the negatives need to keep enough local chemistry to remain possible, while still being conformationally wrong. Random perturbations can often be rejected by a single geometric rule, and docking-derived negatives may instead teach the energy function itself. We therefore used one negative-generation rule: the experimental glycan was moved as a rigid body, poses with 4-8 Å RMSD were kept, and we then selected poses by the number of polar or CH-π contact edges that remained. This rule gives a controlled gradient from low-contact to high-contact negatives, and it does not introduce a separate docking-energy target.

We treated this problem as pocket-conditioned glycan conformation discrimination. First, we defined MIEN, which is an equivariant graph network built on a minimal protein-glycan interface; only protein atoms that form polar or CH-π contacts with the glycan are kept. Cross edges are present only when a semantic contact exists, and protein-protein edges are not allowed. Second, we built one unified negative family by rigidly perturbing experimental complexes and selecting three contact tiers with at least 10 semantic contact edges, 6-9 edges and 3-5 edges, respectively. Third, we tested MIEN with a protein-disjoint split, and the tests included tier-specific discrimination, within-pocket score separation and input ablations. Fourth, we tested MIEN on the 260 held-out positive complexes with candidate poses generated by AutoDock-GPU and GlycanDock. MIEN and the software score ranked the same case-matched pools. We compared the symmetry-corrected RMSD of each docking engine’s best pose with the pose that received the highest MIEN score, and RMSD was not used to generate or filter candidates. Cross-sugar ranking was not tested.

## 2. Methods

### 2.1 Problem formulation and inputs

MIEN is a structure-based equivariant graph network that gives a score for a glycan candidate pose inside a fixed protein pocket. The model returns a scalar compatibility score and ranks different poses of the same glycan; it uses only heavy-atom coordinates from the Protein Data Bank (PDB) ^40^ and local chemistry from the protein-glycan complex. Protein sequence, residue identities, chain identifiers and sugar residue names are not given to the model; for this reason, MIEN can only rank poses inside a known pocket.

### 2.2 Dataset, leakage control and splits

The final dataset contains 6,889 protein-glycan complexes, including 2,634 experimental poses and 4,255 non-native poses. Positive examples were the curated experimental protein-carbohydrate complex PDB files released with CLIMBS^21^. These PDB files were used directly without redocking, chain transfer, coordinate minimization or ligand reconstruction. For model input, protein ATOM records were assigned to the receptor and carbohydrate HETATM residues to the glycan ligand. Hydrogen atoms, water, ions, additives and non-carbohydrate cofactors were not included in the interface graph. The protein and glycan heavy-atom coordinates in the CLIMBS files were kept, no missing atoms were rebuilt, and only protein atoms forming at least one semantic contact with the glycan were retained. All negatives were generated by one contact-preserving perturbation rule. The three semantic-contact tiers contributed 1,460, 1,454 and 1,341 negatives in the final usable dataset.

Partitions follow the similarity-cluster split; all records from one cluster stay in the same partition. The resulting partitions contain 5,241 training rows (2,114 positive and 3,127 negative), 820 validation rows (260 positive and 560 negative) and 828 test rows (260 positive and 568 negative). No test or validation protein shares the similarity-cluster criterion with a training protein.

To separate training discrimination from pose-selection performance, the 260 positive complexes in the test partition were kept for the redocking benchmark. These complexes covered 229 unique proteins and 17 glycan residue types. Protein entries were clustered before splitting. Two entries entered the same cluster when their global sequence similarity was at least 90%, or when a local alignment reached at least 90% similarity over at least 80% of the shorter sequence. Independent realignment of 1,616 proteins found no cross-partition hit under this criterion; the closest cross-partition pair had 89.85% effective similarity and 98.53% coverage. These cases were not used for model fitting, checkpoint selection or score calibration.

### 2.3 Minimal-interface graph

Each complex was turned into an atom-level graph in three steps.

First, a protein atom was kept only when it formed at least one semantic contact with the glycan. A semantic contact is either a polar contact or a CH-π contact, and both need compatible chemistry and geometry. A polar contact needs a polar protein role and a polar ligand role at a distance of 2.4-3.5 Å. A CH-π contact needs an aromatic protein carbon and a ring, anomeric or exocyclic ligand carbon at 3.0-4.2 Å (**Figure 1**). Protein atoms without a semantic contact were removed, so the interface size was limited by chemistry rather than by a fixed radius. The CH-π geometry follows structural and energetic analyses of carbohydrate-aromatic interactions ^41,42^.

**Figure 1.**
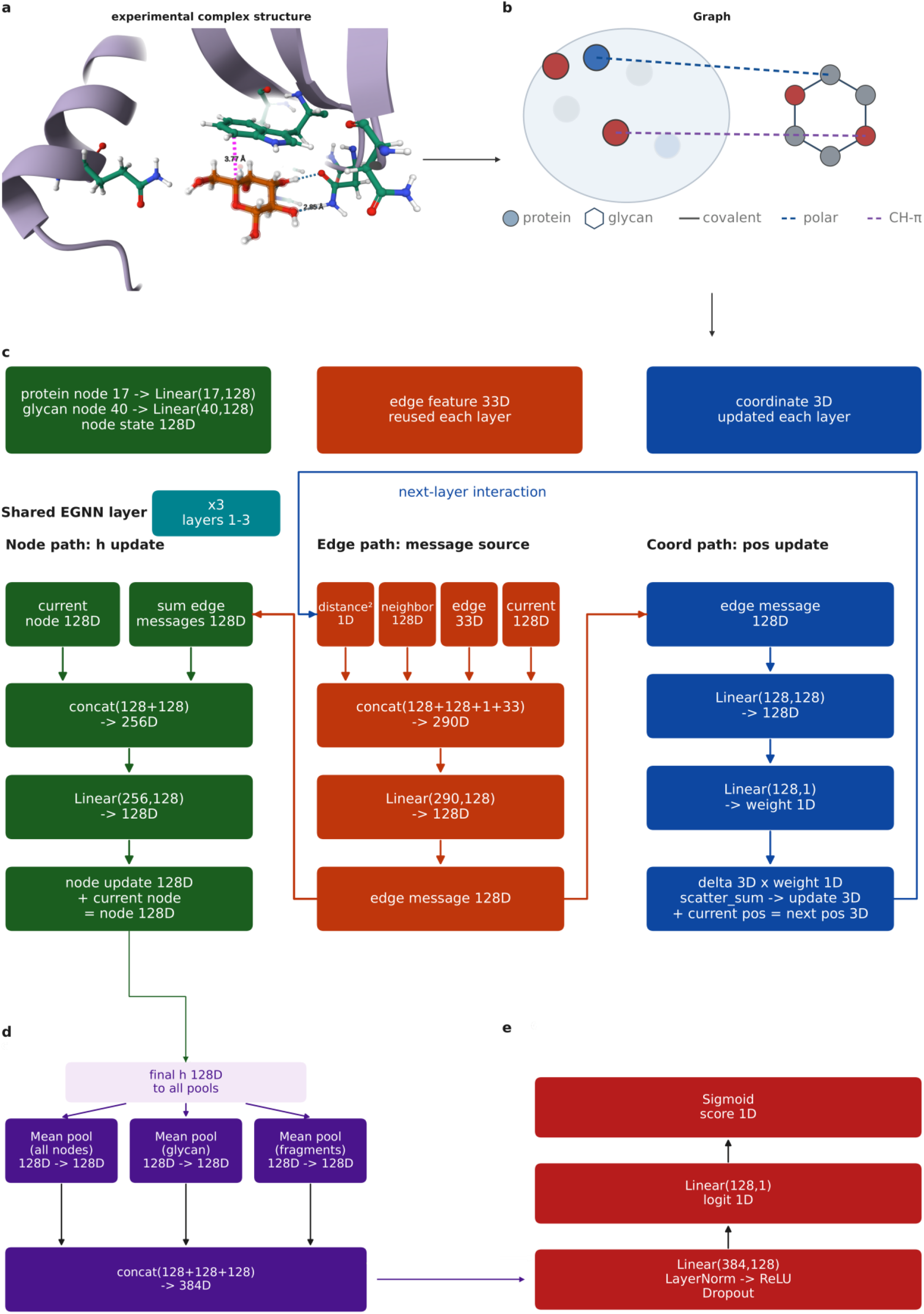
Experimental interface of 1G8Z-GAL and MIEN architecture. a, Protein environment surrounding GAL in PDB 1G8Z. The purple dashed line marks an annotated CH-π contact at 3.77 Å, and the blue dashed line marks a polar contact at 2.85 Å. b, Minimal-interface graph. Only protein atoms forming semantic polar or CH-π contacts are retained; gray edges connect glycan atoms, and blue and purple dashed edges mark polar and CH-π protein-glycan contacts, respectively. c, Three shared EGNN pathway columns for edge, node and coordinate updates. Protein (17-d) and glycan (40-d) node features are projected to 128-d states, whereas edge features remain 33-d. Each MLP block is Linear, LayerNorm, ReLU, Dropout and Linear. d, Mean pooling over all atoms, glycan atoms and glycan fragments gives three 128-d vectors, which are concatenated into a 384-d representation. e, A 384-128-1 MLP produces one logit, and a sigmoid gives the interface-compatibility score.

Second, edges were made along covalent bonds inside the glycan and across every semantic protein-glycan contact. The edges were undirected; no protein-protein edges were made, because protein-internal connectivity carries fold and size information that is not part of the question and could allow protein-similarity leakage. No generic distance contact edge was used, so every cross edge was semantic by construction.

Third, each edge received a 33-dimensional feature vector, which is described in Section 2.4.

Formally, each candidate complex is written as an atom-level graph G=(V,E). The node set is divided into protein nodes VP and glycan nodes VL, and the edge set contains glycan-covalent edges EL and semantic cross edges EPL:

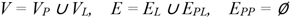

Polar and CH-π contacts are defined by chemistry and distance, not by one distance cutoff:

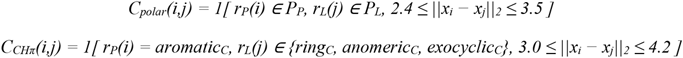

The graph is not meant to be a complete physical model of the complex. It is a controlled interface representation, and it removes protein fold, protein size and generic packing information that could otherwise support identity or dataset shortcuts.

### 2.4 Model architecture

Nodes encode local chemistry but not molecular identity.

Protein atoms receive 17 features, including element type, residue chemical class, a backbone flag and an interaction role. The role definition is the same as the one used in the polar and CH-π contact rules, so node and edge semantics stay consistent. For protein nodes, unreachable sulfur, selenium, hydrogen and metal channels were omitted (**Figure 1c-e**).

Ligand atoms receive 40 features, including element type, sugar role, hybridization, heavy-atom degree, attached-hydrogen count, heteroatom-neighbor count and six chemistry flags. These features describe charged and decorated sugars, but the residue name is not provided.

Edge features are 33-dimensional. They include five bond-type channels, 16 Gaussian distance channels over 0-5 Å, two semantic contact channels, six orientation features, and four glycosidic-torsion channels (sin/cos of phi and psi; zero on non-glycosidic edges). Cross edges carry an all-zero bond block.

Unreachable, duplicated or constant channels were omitted, and they are listed in the feature schema and the released source code; the retained features describe local chemistry and geometry, not protein or glycan identity.

The edge representation is

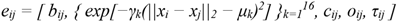

where bij is the covalent bond block, the second term is a 16-channel Gaussian distance expansion, cij contains the polar and CH-π semantic channels, oij contains local orientation terms, and τij contains glycosidic torsion terms and is zero on other edges. Cross edges carry an all-zero bond block.

The encoder follows an EGNN-style equivariant update ^43,44^, and it was adapted so that coordinate updates follow the minimal-interface graph. The equivariance contract is explicit, i.e., for a proper rotation Q ∈ SO(3) and translation t, node coordinates change as xi ↦ Q xi + t, while node states and the scalar interface score stay invariant.

Protein and glycan atoms use separate input projections, and the projections are merged into a shared hidden state:

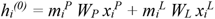

where mi^P^ and mi^L^ select the protein and glycan branches. For layer l, an edge message is formed from both endpoint states, the squared interatomic distance and eij:

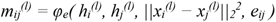

Incoming messages are added together and passed to a node-update MLP:

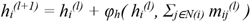

Coordinates are moved along edge directions by a learned scalar bounded by tanh, and the global scale c is initialized near zero:

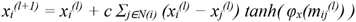

Because the graph has no protein-protein edges, coordinate updates move only atoms that are in protein-glycan or glycan-glycan contacts. The reported model uses three layers, a hidden dimension of 128 and dropout 0.2; no attention mechanism is used.

These design choices come from the decision problem that MIEN is meant to solve. MIEN compares poses of the same glycan inside a fixed pocket, so the score should not change when the whole system is rotated or translated; at the same time, the score should still respond to directional local geometry. The equivariant update meets this need because it updates node states and coordinates together. The graph keeps only semantic cross edges, because protein-internal connectivity would bring back protein fold and size information. Those features are not part of the local compatibility question, and they could create an identity-based shortcut. We use pairwise edge messages instead of global attention. Each message already depends on both endpoint states, the squared distance and the chemical edge type; global attention would bring back the long-range protein context that the minimal-interface construction is meant to remove. Pilot training runs with more message-passing layers did not improve validation performance, so the encoder was fixed at three layers. The 128-dimensional hidden state and dropout of 0.2 were kept as compact regularization settings, and we do not claim that they are globally optimal. The effects of the retained edge, node and coordinate-update components are tested by frozen-checkpoint ablations in Section 3.2.

After message passing, three graph summaries are concatenated including the mean over all retained atoms, the mean over glycan atoms and the mean over glycan fragments:

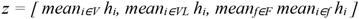

The first vector summarizes the local interface, the second summarizes the whole candidate glycan, and the third summarizes its residue-level organization. A two-layer MLP maps z to one logit, and the sigmoid output is reported as an interface-compatibility score:

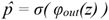

This score is not a calibrated binding probability, an affinity estimate or a general confidence measure; it is only meant for comparing candidate conformations of the same glycan inside the same pocket under the evaluation protocol in Section 2.7.

### 2.5 Negative construction

Negative examples were generated from the experimental protein-glycan complex, and no docking energy was used (**Figure 2**). One contact-preserving perturbation rule was applied to each unique protein-glycan group; the glycan was rotated around its heavy-atom centroid and translated by up to 1.5 Å in each coordinate direction. A candidate pose was kept only if the symmetry-corrected heavy-atom RMSD was between 4 and 8 Å, the minimum protein-glycan heavy-atom distance was at least 1.5 Å, at least two different protein atoms formed polar or CH-π contacts, and at least three semantic contact edges were present.

**Figure 2.**
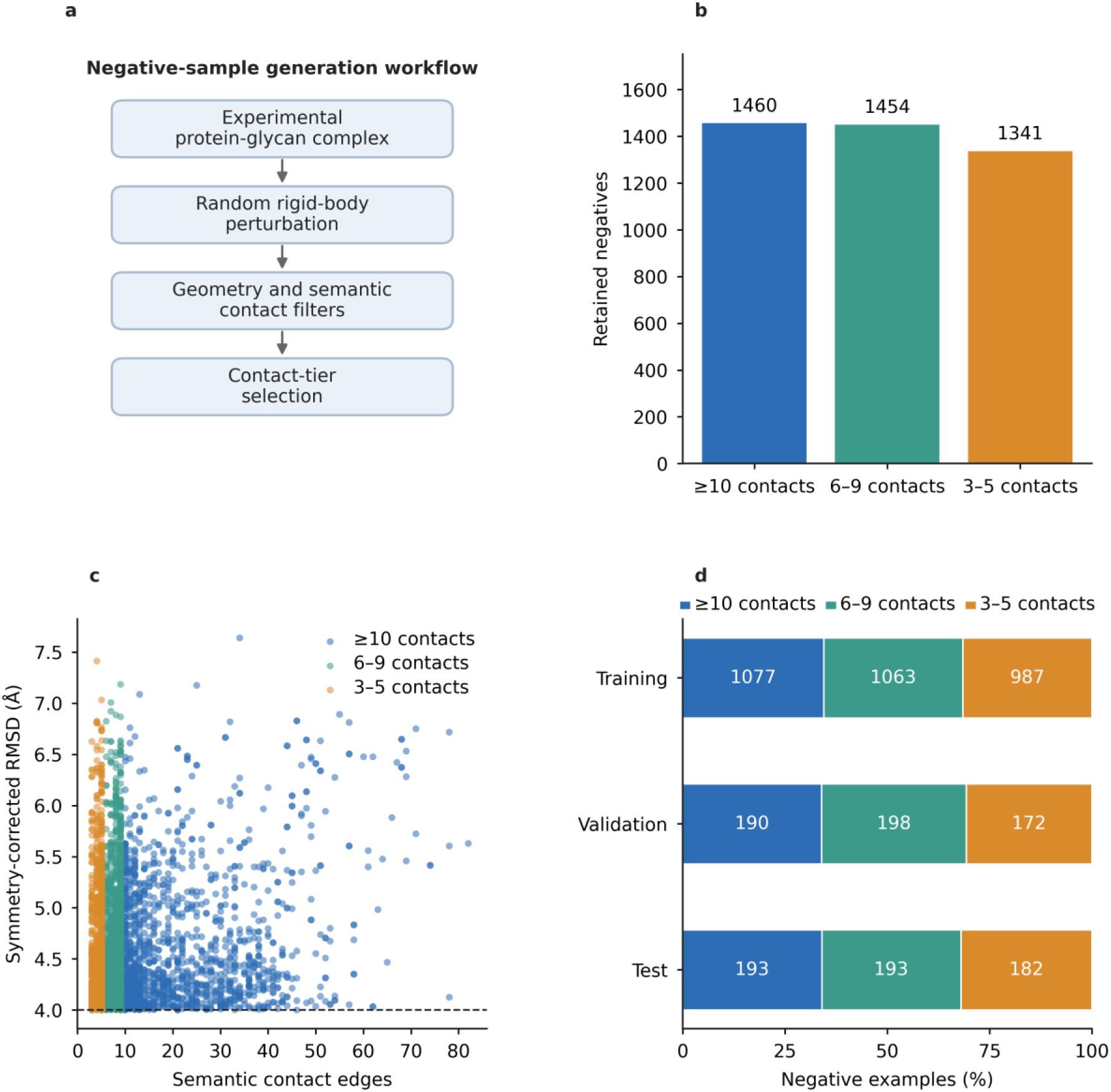
Construction of contact-preserving negatives. a, Vertical workflow for negative generation, from experimental complexes through rigid-body perturbation and semantic-contact filtering to contact-tier selection. b, Number of retained negatives in the >=10, 6-9 and 3-5 semantic-contact tiers. c, Distribution of semantic contact edges and symmetry-corrected RMSD; the dashed line marks 4 Å. d, Contact-tier composition of the training, validation and test partitions, with bar labels showing negative counts. Panels b-d use the 4,255 graph-compatible negatives retained for model training.

The accepted poses were divided into three contact tiers. The highest tier required at least 10 semantic contact edges, the middle tier required 6-9 edges, and the lowest tier required 3-5 edges. For each protein-glycan group, at most one pose was kept in each tier, so there were at most three negatives per group.

Complete complexes were deduplicated by content hash; this design keeps all negatives in the same pocket context and prevents a simple contact-loss shortcut.

The final generation run produced 4,268 contact-preserving negatives before graph construction and 4,255 usable negatives after filtering. The usable tiers contained 1,460 poses with at least 10 edges, 1,454 poses with 6-9 edges and 1,341 poses with 3-5 edges. RMSD and contact counts were kept as metadata, but they were not supplied to the model.

### 2.6 Training

Models were trained with binary cross-entropy, the AdamW optimizer and a learning rate of 3 x 10^-5^. The other settings were weight decay 1 x 10^-4^, batch size 32, dropout 0.2, hidden dimension 128 and three message-passing layers. Model selection used validation AUC, with early stopping after 20 epochs without improvement and no fixed epoch limit. We trained three seeds (42, 43 and 44). The selected epochs were 73, 124 and 134, and training stopped after 93, 144 and 154 epochs, respectively. Training used 5,241 rows and 820 validation rows. Across the three seeds, validation AUC at the selected checkpoints was 0.981, 0.984 and 0.985.

### 2.7 Evaluation protocols

#### Protein-split discrimination

All three partitions were evaluated with AUC, AUPR, accuracy at a score threshold of 0.5, balanced accuracy, sensitivity, specificity and MCC. Records from one protein were kept in a single partition. For the unbalanced test set, AUC, AUPR, MCC and balanced accuracy were given more weight than accuracy alone.

#### Within-pocket and contact-tier discrimination

The inferential analysis used protein-glycan groups that contained at least one experimental pose and at least one contact-preserving negative. For the >=10, 6-9 and 3-5 edge tiers, we computed ROC AUC, AUPR and MCC against the matched positives. Pockets without a same-pocket negative in a tier were excluded only from that tier-specific analysis.

#### Contact-tier stratification

All negatives use the same generation rule, and they differ only in the number of retained polar or CH-π contact edges. Higher-contact negatives are therefore difficult controls, because they keep much of the local interface geometry while staying more than 4 Å from the experimental conformation.

#### Unified held-out docking benchmark

The docking evaluation used all 260 positive complexes in the protein-disjoint test partition. Candidate poses were generated independently with AutoDock-GPU and GlycanDock. The full docking protocols, candidate definitions, ranking endpoints and case-level comparisons are described in Section 4. No case was removed because of MIEN score, docking score, glycan type or RMSD.

### 2.8 Ablation, implementation, and reproducibility

All ablations used the frozen checkpoints from the three seeds and the same 828 held-out test complexes as the discrimination analysis. No parameters were updated. Feature-block ablations zeroed selected protein-node, glycan-node or edge-feature channels before the forward pass. Layer ablations replaced one message-passing layer at a time with an identity operator, while the other layers retained their parameters and graph topology. The coordinate-update ablation set the learned coordinate scale of every layer to zero. We report AUC and MCC for each ablated model and their decreases relative to the full model. Results are summarized as mean ± s.d. across seeds (**Tables 2 and 3**; **Figure 3c-f**).

**Figure 3.**
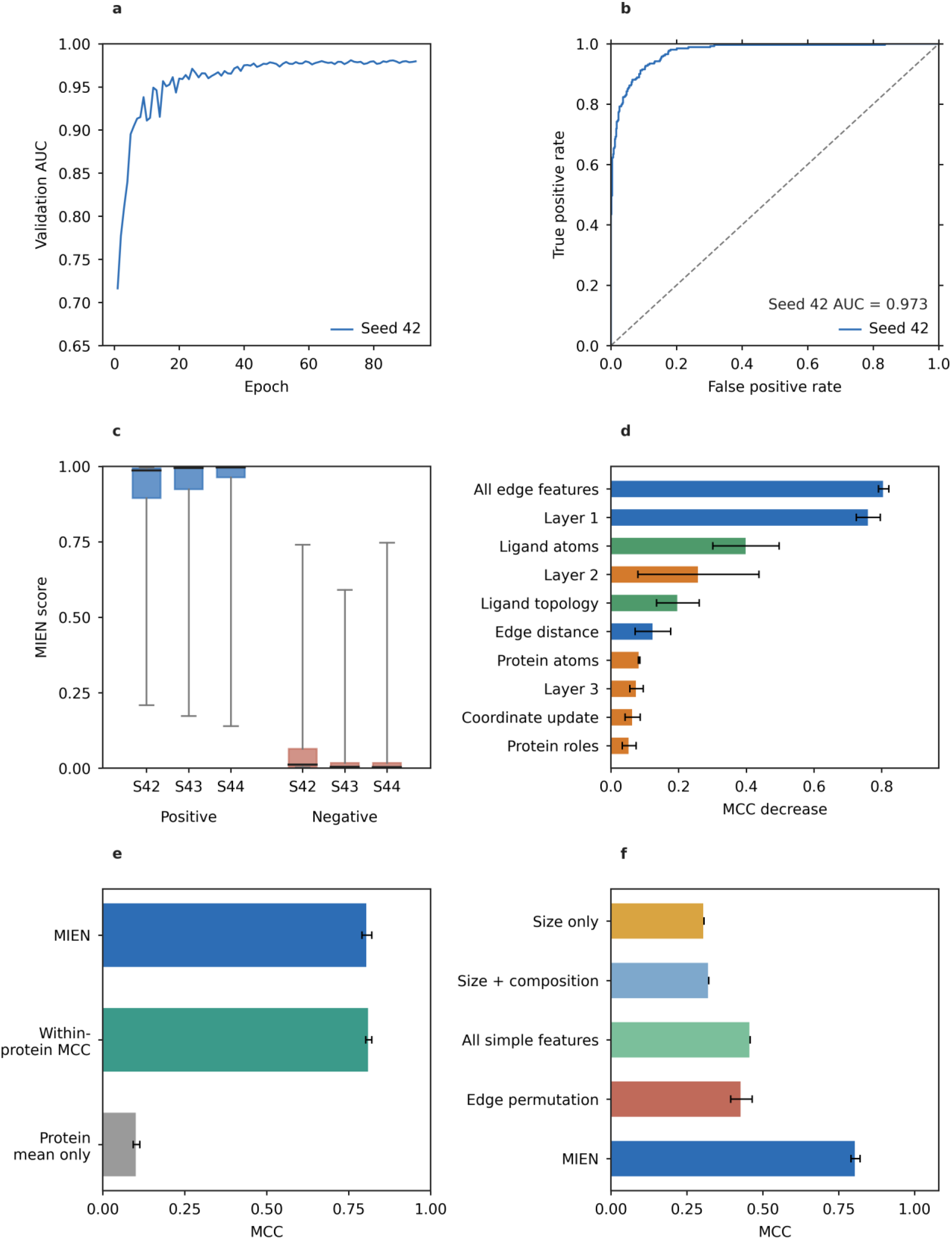
Discrimination and ablation of MIEN trained on the unified negative set. a, Validation AUC curve for seed 42. b, ROC curve for seed 42 on the held-out test partition. c, Distributions of MIEN scores for experimental positives and contact-preserving negatives across three seeds; boxes show the median and interquartile range, and whiskers extend to the 5th and 95th percentiles. d, Feature and layer ablations ordered by mean MCC decrease, with error bars showing the standard deviation across three seeds. e, Overall MIEN MCC, within-protein macro MCC and protein-mean-only MCC across three seeds. f, MCC of interface-summary baselines and cross-edge feature permutation, compared with MIEN across three seeds.

Graphs were built once and cached, and the cache was invalidated when the feature schema changed. Models were implemented in PyTorch and PyTorch Geometric ^45^. Ligand chemistry (including bond orders, ring membership, hydrogen counts, carbonyl and carboxylate detection) used RDKit ^46^, with a purely geometric fallback when the deposited file was incomplete. Docking used AutoDock-GPU ^8^. Every training run and downstream evaluation appended a long-format record to a result registry. Each record contained the checkpoint path, checkpoint identifier, dataset, split, dataset size and code version. The tables and figures in this manuscript were assembled from that registry.

## 3. Results

We first tested whether MIEN separated experimental from non-native poses on held-out proteins. We then examined which representation blocks drove that decision and whether simpler interface descriptors could explain the score. Section 4 reports the held-out docking validation, in which MIEN and the corresponding software score ranked case-matched AutoDock and GlycanDock candidate pools.

### 3.1 A unified contact-preserving negative set supports discrimination

The final dataset contained 6,889 protein-glycan complexes, with 2,634 experimental poses and 4,255 contact-preserving negatives. The three contact tiers contributed 1,460 high-contact (>=10 edges), 1,454 middle-contact (6-9 edges) and 1,341 lower-contact (3-5 edges) negatives. Partitions followed the similarity-cluster split. The training partition contained 2,114 positive and 3,127 negative poses, the validation partition contained 260 positive and 560 negative poses, and the test partition contained 260 positive and 568 negative poses. The distributions are summarized (**Table 1** and **Figure 2**).

**Table 1.** Training partitions for the unified negative-generation rule.

| Partition | Positive | Contact-preserving | Total |
| --- | --- | --- | --- |
| Training | 2,114 | 3,127 | 5,241 |
| Validation | 260 | 560 | 820 |
| Test | 260 | 568 | 828 |
| Total | 2,634 | 4,255 | 6,889 |

On the 828-complex test partition, the three seeds reached AUC 0.974 ± 0.002, AUPR 0.952 ± 0.003, MCC 0.806 ± 0.015, accuracy 0.914 ± 0.008, balanced accuracy 0.911 ± 0.005, sensitivity 0.904 ± 0.007 and specificity 0.918 ± 0.013 (**Table 2**; **Figure 3**). Because the negatives retain semantic contacts, AUC, AUPR and MCC were emphasized over accuracy alone.

**Table 2.** Held-out discrimination of the model trained on the unified negative set.

| Metric | Value |
| --- | --- |
| AUC | $0.974 \pm 0.002$ |
| AUPR | $0.952 \pm 0.003$ |
| Accuracy at 0.5 | $0.914 \pm 0.008$ |
| Balanced accuracy at 0.5 | $0.911 \pm 0.005$ |
| Sensitivity at 0.5 | $0.904 \pm 0.007$ |
| Specificity at 0.5 | $0.918 \pm 0.013$ |
| MCC at 0.5 | $0.806 \pm 0.015$ |

### 3.2 Edge geometry and ligand chemistry carry most of the scoring signal

To identify which parts of the minimal-interface representation supported discrimination, we performed inference-time ablations on the three frozen checkpoints using the same 828 held-out complexes. In the feature-block ablations, selected input channels were set to zero before the forward pass. No model was retrained. We report the mean and standard deviation across the three seeds.

The edge representation was the strongest source of scoring signal. Removing all edge features reduced AUC from 0.974 ± 0.002 to 0.573 ± 0.039 and reduced MCC from 0.806 ± 0.015 to zero. Removing the Gaussian distance channels reduced AUC to 0.936 ± 0.011 and MCC to 0.682 ± 0.040. These results show that edge geometry is needed for the score.

Glycan node features were the next major dependency. Removing all ligand-atom features reduced AUC to 0.833 ± 0.023 and MCC to 0.407 ± 0.092. Removing ligand topology reduced AUC to 0.917 ± 0.009 and MCC to 0.608 ± 0.067. Removing protein-atom features reduced AUC to 0.942 ± 0.006 and MCC to 0.722 ± 0.012. Protein element, group and role ablations produced smaller decreases, which indicates that local protein chemistry still contributes to the decision.

All edge features were the strongest dependency, followed by the first EGNN layer, ligand atoms, the second layer, ligand topology and edge distance. Removing the first message-passing layer reduced AUC to 0.743 ± 0.054 and MCC to 0.045 ± 0.048. The coordinate-update ablation reduced AUC to 0.962 ± 0.003 and MCC to 0.742 ± 0.009, so coordinate motion contributed to the score but was less important than edge and ligand features. The within-protein macro MCC was 0.812 ± 0.009 across 221 informative pockets, whereas a protein-mean-only score reached MCC 0.103 ± 0.010. Logistic baselines using interface size alone, size plus composition, and size plus composition plus aggregate contacts reached MCC 0.307, 0.323 and 0.458, respectively, below MIEN at MCC 0.806 ± 0.015. Permuting cross-edge feature vectors while preserving graph topology decreased MCC to 0.430 ± 0.035 across five permutations per seed. These controls support a within-pocket geometric component and show that the model is not simply a protein-similarity or interface-size classifier (**Table 3** and **Figure 3d-f**).

**Table 3.** Feature, layer and coordinate ablations of the model trained on the unified negative set, ordered by MCC decrease.

| Ablation | AUC | MCC | MCC decrease |
| --- | --- | --- | --- |
| Baseline | 0.974 ± 0.002 | 0.806 ± 0.015 | 0.000 |
| All edge features | 0.573 ± 0.039 | 0.000 ± 0.000 | 0.806 |
| Layer 1 | 0.743 ± 0.054 | 0.045 ± 0.048 | 0.761 |
| Ligand atoms | 0.833 ± 0.023 | 0.407 ± 0.092 | 0.399 |
| Layer 2 | 0.929 ± 0.039 | 0.547 ± 0.187 | 0.259 |
| Ligand topology | 0.917 ± 0.009 | 0.608 ± 0.067 | 0.198 |
| Edge distance | 0.936 ± 0.011 | 0.682 ± 0.040 | 0.124 |
| Protein atoms | 0.942 ± 0.006 | 0.722 ± 0.012 | 0.084 |
| Layer 3 | 0.955 ± 0.008 | 0.731 ± 0.005 | 0.075 |
| Coordinate update | 0.962 ± 0.003 | 0.742 ± 0.009 | 0.064 |
| Protein roles | 0.964 ± 0.005 | 0.752 ± 0.031 | 0.054 |
| Ligand roles | 0.961 ± 0.009 | 0.760 ± 0.014 | 0.046 |
| Ligand elements | 0.970 ± 0.001 | 0.773 ± 0.009 | 0.033 |
| Protein elements | 0.965 ± 0.002 | 0.775 ± 0.007 | 0.031 |
| Protein groups | 0.969 ± 0.000 | 0.788 ± 0.005 | 0.018 |

## 4. Unified held-out docking validation

### 4.1 Benchmark construction and docking protocol

The held-out docking benchmark was built from all 260 positive complexes in the test partition. These complexes contained 229 unique proteins and 17 glycan residue types. Protein sequences were clustered at 90% similarity with 80% coverage of the shorter sequence before the split. No case was selected according to MIEN score, AutoDock score, glycan class or RMSD; the experimental glycan was removed from the receptor, the curated protein coordinate set was kept, and the same glycan was docked into its original pocket.

Both engines were run with their official recommended protocol settings, and each engine generated 20 candidate poses per case. AutoDock-GPU used 20 independent LGA searches per complex, ligand-size-based heuristic search, automatic stopping, 2,500,000 energy evaluations per search and the default 12,000,000 heuristic asymptotic cap. One entity-1 pose was kept from each LGA search. Receptor grids and ligand parameters were prepared once per case. The experimental pose, MIEN score and RMSD were not used to generate, rank or filter the AutoDock candidates.

The same complexes were docked with PyRosetta GlycanDock. The official benchmark-recommended GlycanDock settings were used including three repeats, eight rigid-body rounds and eight torsion rounds. Twenty independent trajectories were generated per complex to match the AutoDock candidate-pool size, and seed 42 was used. The receptor and glycan were taken from the experimental complex. The glycan was mapped to its Rosetta carbohydrate type and terminal patches. GlycanDock scoring used the recommended carbohydrate-specific weights in the ref2015 energy function (*sugar_bb* = 0.5; *fa_intra_rep_nonprotein* = 0.55). No GlycanDock or Rosetta score was given to MIEN. RMSD was evaluated in the receptor frame over the connected heavy-atom core shared by the mapped native pose and each docked trajectory. Poses without a semantic protein-glycan interface did not receive a MIEN score. The GlycanDock-selected pose was the trajectory with the lowest Rosetta interaction_energy.

### 4.2 Unified best pose comparison across docking engines

AutoDock and GlycanDock each generated 20 candidate poses per case. The corresponding software score and the seed-42 MIEN checkpoint ranked the same case-matched candidate pools, and symmetry-corrected RMSD was calculated only after the ranking was fixed. The analysis compared the best pose selected by the docking engine with the highest score pose selected by MIEN. Top-k aggregates were not used, because RMSD values from different cases are not directly comparable; for GlycanDock, the software ranking used all valid trajectories, while MIEN ranked only poses for which a minimal interface could be constructed.

Across the held-out cases, MIEN selected lower-RMSD best poses in 147 of 260 AutoDock cases and 112 of 260 GlycanDock cases, while the software selected lower-RMSD poses in 83 and 88 cases, respectively. The other cases were ties. MIEN also increased the number of cases below 1, 2 and 4 Å for both engines. For AutoDock, the counts increased from 33 to 37, 43 to 53 and 81 to 97; for GlycanDock, they increased from 18 to 38, 35 to 74 and 79 to 105. The paired differences and threshold counts are summarized (**Figure 4**).

**Figure 4.**
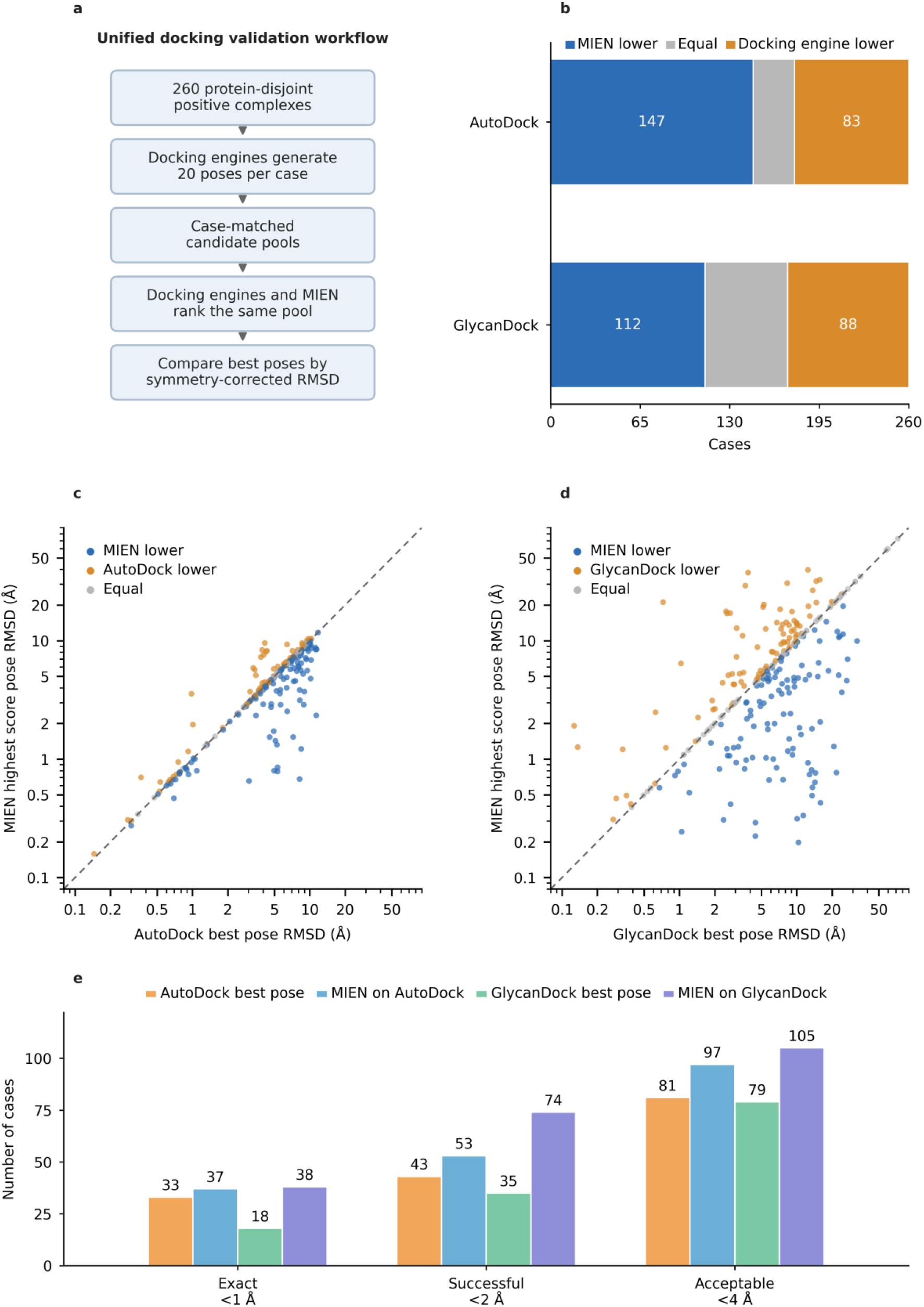
Unified AutoDock and GlycanDock validation of MIEN trained on the unified negative set. The seed-42 checkpoint was used for reranking. a, Unified docking validation workflow. b, Case-level counts of lower-RMSD selections. c,d, Paired AutoDock and GlycanDock RMSDs; blue points indicate a lower MIEN RMSD, orange points a lower software RMSD and gray points ties. e, Cases below 1, 2 and 4 Å. n=260 held-out complexes.

### 4.3 Glycan-dependent performance and failure boundary

The case-level comparison showed a boundary that depended on the glycan and the engine. No single glycan explained the residual failures; cases in which the docking engine selected the lower-RMSD pose were spread across multiple sugar types. Performance depended on the candidate engine, pocket geometry and the specific pose, and glycan types with few held-out cases are interpreted with caution. Candidate-pool composition also places a ceiling on reranking (Table 4).

**Table 4.**
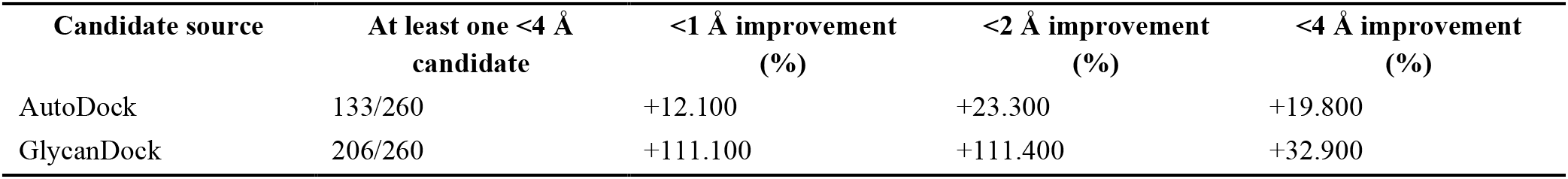
Candidate-pool ceiling and RMSD-threshold improvements. Relative improvement is calculated as (MIEN highest score pose - docking engine best pose) / docking engine best pose.

| Candidate source | At least one <4 Å candidate | <1 Å improvement (%) | <2 Å improvement (%) | <4 Å improvement (%) |
| --- | --- | --- | --- | --- |
| AutoDock | 133/260 | +12.100 | +23.300 | +19.800 |
| GlycanDock | 206/260 | +111.100 | +111.400 | +32.900 |

Several limitations bound this conclusion. The held-out set contains 260 complexes and one 20-run AutoDock-GPU search set plus one 20-trajectory GlycanDock set per complex. It measures ranking inside these candidate pools, not candidate generation. All 17 glycan types appeared in training, so the engine-dependent patterns reflect pocket and pose context rather than completely unseen sugar classes. The analysis does not establish polysaccharide, multi-glycan, cross-sugar, affinity or binding-site prediction. Candidate-pool composition also places a ceiling on reranking. The candidate-pool results (**Table 4**) summarize whether each engine pool contained at least one sub-4 Å candidate and the relative improvement in sub-1, sub-2 and sub-4 Å selections after MIEN reranking. AutoDock pools contained at least one sub-4 Å candidate in 133 of 260 cases, with improvements of 12.1%, 23.3% and 19.8% at the 1, 2 and 4 Å thresholds. GlycanDock pools contained at least one sub-4 Å candidate in 206 of 260 cases, with improvements of 111.1%, 111.4% and 32.9%, respectively. MIEN is a post-docking reranking model rather than a replacement for energy-based docking scoring, and it does not generate new poses. Its benefit is therefore limited by the quality of the original docking ensemble. In general, a better candidate pool gives MIEN more opportunity to identify a near-native pose. When all candidates are far from the experimental conformation, reranking cannot recover a pose that the docking engines did not generate.

MIEN has no explicit energy term. Its complementary gain over AutoDock and GlycanDock indicates that local interface geometry contains ranking information that is not fully represented by either energy score, but this does not make energy scoring unnecessary. MIEN should be used as a reranking layer together with docking; redocking can also change internal ligand geometry, so this benchmark does not separate rigid placement errors from internal torsional or ring-conformation errors.

## 5. Discussion

The main finding is that local protein-glycan geometry supports a bounded but useful decision about which glycan state belongs in a pocket. MIEN trained on the unified negative set achieved AUC 0.974 ± 0.002, AUPR 0.952 ± 0.003 and MCC 0.806 ± 0.015 across three seeds on the held-out discrimination test. In the unified docking benchmark, MIEN selected lower-RMSD top-1 poses in 147 of 260 AutoDock cases and 112 of 260 GlycanDock cases, and it increased the number of high-accuracy best poses. The benefit still depended on the docking engine, pocket geometry and glycan type.

MIEN addresses a narrower question than the models described above. Given a fixed pocket and a fixed glycan, it ranks candidate poses by local interface compatibility. It does not predict binding sites, affinity or cross-sugar preference. This scope makes the comparison with docking scores direct, because all methods rank the same candidate pool.

Three-seed training produced consistent discrimination performance. The within-protein macro MCC and shortcut controls show that the score is not driven by protein identity, interface size or global composition. Edge geometry and glycan atom chemistry were the main dependencies; coordinate updates contributed to the score but were less important than the edge and ligand blocks.

The negative construction matters for that claim. All negatives followed the same rigid-perturbation rule, and they differed only in how many semantic contact edges were retained. The ablation results show that the score depends strongly on edge geometry and glycan atom features, and the shortcut controls show that simple protein-mean and interface-size descriptors do not reproduce the result.

Because MIEN and the software score saw the same case-matched candidate pools, the comparison should be read as candidate reranking, not as a replacement for docking. MIEN improved top-1 selection for both engines, but the gains were not uniform across proteins, glycans or docking engines; local interface geometry adds information beyond the two energy scores, while the software rankings remain necessary for candidate generation and for many accurate best pose selections.

Several boundaries constrain the result. The benchmark contains 260 held-out complexes and one 20-run AutoDock-GPU search set plus one 20-trajectory GlycanDock set per complex. It supports best pose ranking inside these candidate pools, but it does not establish candidate generation, polysaccharide or multi-glycan docking, or cross-sugar ranking. All glycan residue types appeared in training, so the observed differences are related to specific protein pockets and poses, not to a completely unseen sugar class; MIEN also has no explicit energy term, so it should complement energy-based docking scores instead of replacing them.

The docking analysis shows how MIEN should be used. It improved threshold success more strongly for GlycanDock than for AutoDock, while AutoDock had more case-level top-1 wins. This difference follows from candidate-pool quality, i.e., reranking can only recover a near-native pose if that pose is already present. MIEN is therefore best used as a post-docking reranking layer rather than a standalone pose generator.

Two follow-up studies would address the most important remaining uncertainties. First, a conformation-controlled decoy suite should hold protein-glycan placement fixed while varying internal torsions or ring states; this would test whether the discriminator reads conformational compatibility instead of only placement. Second, extending MIEN from monosaccharides to polysaccharide and multi-glycan docking should explicitly model glycosidic linkage flexibility, multiple glycan chains and their coupled placement inside the pocket.

## 6. Code availability

The MIEN implementation required for graph construction, training and pose scoring is available at https://github.com/Huangxirui666/MIEN.

